# Treenome Browser: co-visualization of enormous phylogenies and millions of genomes

**DOI:** 10.1101/2022.09.28.509985

**Authors:** Alexander M. Kramer, Theo Sanderson, Russell Corbett-Detig

## Abstract

Treenome Browser is a web browser tool to interactively visualize millions of genomes alongside huge phylogenetic trees.

**Availability and Implementation:** Treenome Browser for SARS-CoV-2 can be accessed at cov2tree.org, or at taxonium.org for user-provided trees. Source code and documentation are available at github.com/theosanderson/taxonium and docs.taxonium.org/en/latest/treenome.html.

## Introduction

Huge genomic datasets are increasingly common and widely available. The ongoing COVID-19 pandemic has particularly stressed existing bioinformatic infrastructure for analysis of genomic data. We developed UShER (Turakhia *et al*., 2021) to enable the inference of a continuously growing phylogeny containing millions of SARS-CoV-2 genomes (McBroome *et al*., 2021), and Taxonium to enable interactive exploration of phylogenies of this size (Sanderson, 2022). Gingr (Treangen *et al*., 2014) and the UCSC Genome Browser can display a multiple sequence alignment alongside a phylogeny (Lee et al. 2022). However, no existing tool can simultaneously visualize million sample phylogenies with their underlying genome sequences.

We present Treenome Browser, a visualization tool for exploring genetic variation in millions of genomes alongside a huge phylogeny. Treenome Browser uses an innovative phylogenetic compression technique to interactively display the genome of each sample aligned with its phylogenetic position, remaining performant on trees with over 12 million sequences. The web application is available at cov2tree.org for the global SARS-CoV-2 tree and at taxonium.org for user-provided trees.

## Description

Treenome Browser displays mutations as vertical lines spanning the mutation’s presence among samples in the phylogeny, drawn at their horizontal position in an associated reference genome (Fig. 1). Both amino acid and nucleotide mutations are supported. When the whole tree is visible, the genetic signatures of major clades in the tree become apparent. Users can search or manually zoom to samples in the tree to inspect smaller clades and individual genomes. The genome alignment can also be navigated to analyze sites or regions of interest. Potential applications of Treenome Browser include variation-informed primer design, uncovering technical artifacts in tree construction or primary sequence assembly, and investigating molecular evolutionary processes.

**Figure 1.**
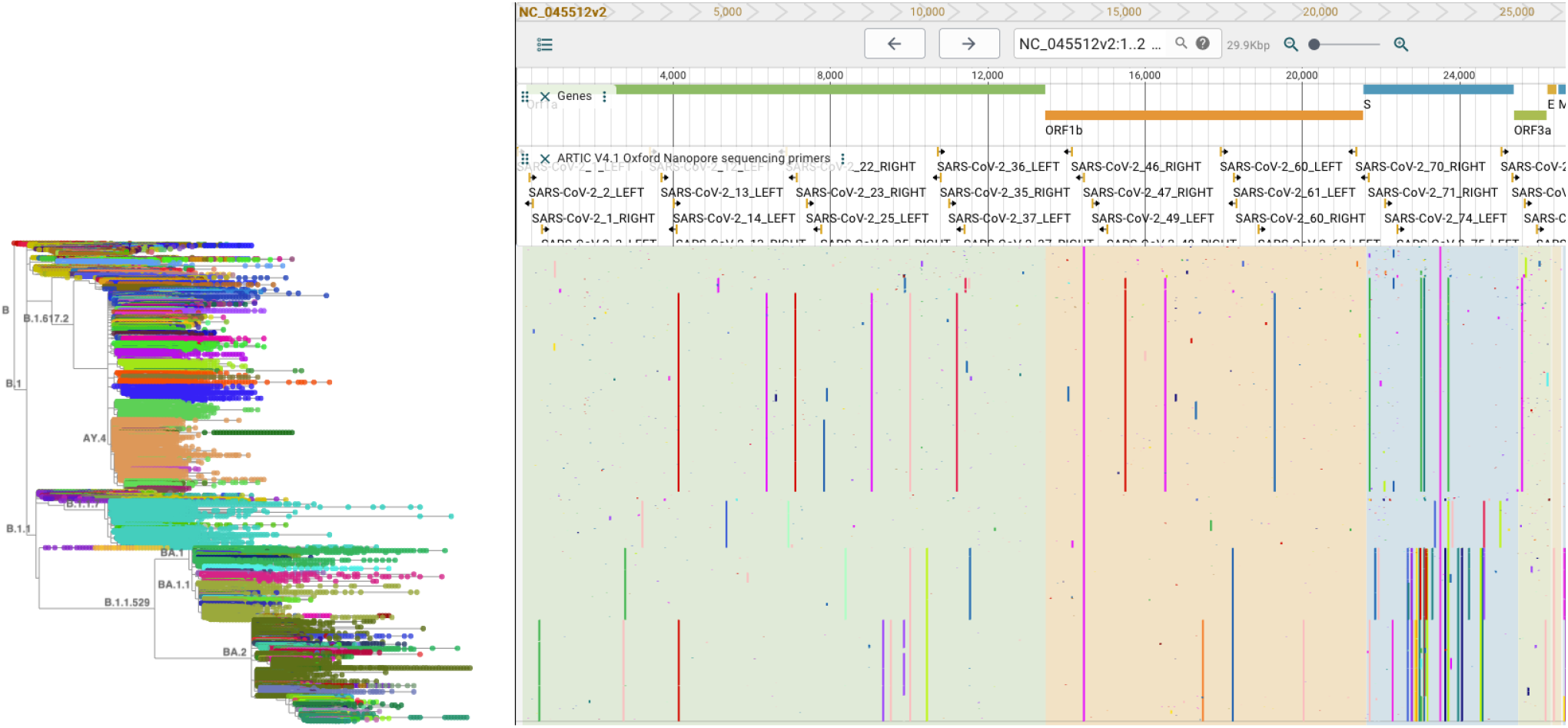
Treenome Browser for the global SARS-CoV-2 tree with 5.9 million genomes, shown with the ARTIC Primers v4.1 (Scalene, 2021) annotation track. Colored lines represent amino acid mutations relative to the reference genome.

We provide a special build of Treenome Browser for the global SARS-CoV-2 tree at cov2tree.org. This site is preloaded with annotations for the reference genome (RefSeq NC_045512v2) curated by the UCSC Genome Browser (Lee et al. 2022). We also support user-provided mutation-annotated trees created with UShER, which can be produced using the TaxoniumTools package (Sanderson 2022) and loaded at taxonium.org.

## Implementation

The mutation-annotated tree data structure enables Treenome Browser to scale to huge trees. Briefly, this structure encodes genomic data in the form of a phylogeny with the inferred emergence of mutations annotated on each node. The genotype data encoded in a mutation-annotated tree is often many times smaller than the same data in VCF format (Table S1). We use this phylogenetic compression to efficiently display mutations. Treenome Browser is much faster than an existing tool with similar functionality, and scales to much larger phylogenomic datasets (Table S2). For a given mutation-annotated tree, the vertical span of each subclade of the tree is first stored using dynamic programming. The tree is then traversed from root to leaves. As each node is visited, its mutations are drawn across the pre-computed vertical span of its descendant clade. We use the deck.gl framework (Uber, 2016) to render the mutations along the genome, which exploits WebGL to render large datasets much more quickly than traditional approaches for web visualization. This phylogenetically-informed process displays on screen the complete set of amino acid and/or nucleotide substitutions in every leaf genome without explicitly enumerating the mutations in each genome, increasing rendering speed and reducing memory usage (Table S3).

Treenome Browser further increases performance by subsampling nodes and mutations. Because many nodes occupy essentially the same spatial position when the tree is zoomed out, Taxonium dynamically displays sparsified trees from the full tree (Sanderson 2022). Treenome Browser also uses these subsampled trees when displaying mutations to avoid unneeded computation. This tool further filters mutations by discarding those spanning very small clades at a given zoom level. These subsampling steps exclude imperceptible data based on zoom level while allowing users to view any individual node or mutation.

The reference genome and annotation tracks are displayed in an interactive JBrowse 2 (Diesh *et al*., 2022) panel. Annotations are loaded into Treenome Browser automatically for the global SARS-CoV-2 tree at cov2tree.org. We use the UCSC Genome Browser API to fetch each bigBed and bigWig annotation file curated by UCSC for SARS-CoV-2 (Fernandes *et al*., 2020). These annotations and user-provided annotation files are displayed by JBrowse 2.

Treenome Browser is implemented in React as a feature addition to the Taxonium project. Source code is available at https://github.com/theosanderson/taxonium/tree/master/taxonium_web_client.

## Acknowledgements

The authors gratefully acknowledge the many developers of the open-source projects enabling the development of Treenome Browser. We thank Jakob McBroome, Angie Hinrichs, and Yatish Turakhia for their feedback and helpful discussions.

