## Supplementary Data for "Treenome Browser: co-visualization of enormous phylogenies and millions of genomes"

### Table S1

| File | Size (GB) |
|---|---|
| public-2022-09-21.all.masked.pb | 0.5 |
| public-2022-09-21.all.masked.jsonl | 1.1 |
| public-2022-09-21.all.masked.nwk | 0.3 |
| public-2022-09-21.all.masked.vcf | 341 |

We downloaded .pb, .nwk, and .vcf files from http://hgdownload.soe.ucsc.edu/goldenPath/wuhCor1/UShER_SARS-CoV-2/2022/09/21/, representing the global SARS-CoV-2 phylogeny as of 9/21/2022 with 6.2M samples. The .pb file is the UShER-format mutation-annotated tree (MAT). We converted the UShER MAT into a Taxonium MAT (*public-2022-09-21.all.masked.jsonl*) with TaxoniumTools. Both the UShER MAT and the Taxonium MAT store a phylogenetic tree and genotype data for that tree, encoding essentially the same information as the combination of a Newick and VCF file in over 300 times smaller size. The Taxonium MAT also stores sample metadata and lineage assignments.

### Table S2

| # samples | Gingr memory usage (GB) | Taxonium + Treenome web browser memory usage (GB) | Gingr loading time (s) | Taxonium + Treenome loading time (s) |
|---|---|---|---|---|
| 5,000 | 1.32 | 0.32 | 4.6 | 2.4 |
| 50,000 | 12.99 | 0.57 | 34.3 | 3.5 |
| 500,000 | | 0.92 | | 8.8 |
| 5,000,000 | | 1.4 | | 25.4 |

We measured the time taken and memory used by Gingr v1.3 and Taxonium + Treenome Browser to load multiple sequence alignments of increasing size. We extracted pruned mutation-annotated tree (MAT) and VCF files from the 9/21/2022 global SARS-CoV-2 phylogeny (https://hgdownload.soe.ucsc.edu/goldenPath/wuhCor1/UShER_SARS-CoV-2/). We then converted the VCF files to compressed Gingr files with HarvestTools and converted the MAT files to Taxonium format with TaxoniumTools. We excluded phylogenetic trees from Gingr analyses because we could not successfully convert trees with over ~10k samples and Gingr crashed when loading large Newick files directly. Cells in gray failed to load (crashed). We ran Gingr on an Ubuntu machine with 126 GB of RAM and loaded the Taxonium trees at taxonium.org on Google Chrome. Peak memory usage was measured with the GNU *time* program for Gingr and the Chrome Task Manager for Taxonium + Treenome Browser. We note that these results are for the client-side version of Taxonium + Treenome Browser and that the server-side version at cov2tree.org loads ~6M samples in about 5 seconds.

**Table S3**

**A (client-side Taxonium)**

| # nodes | Naive algorithm compute time (ms) | Treenome algorithm compute time (ms) | Naive algorithm time to render (s) | Treenome algorithm time to render (s) | Naive algorithm web browser memory usage (MB) | Treenome algorithm web browser memory usage (MB) |
|---|---|---|---|---|---|---|
| 18,921 | 694 | 32 | 2.9 | 2.4 | 453 | 102 |
| 58,253 | 1,762 | 103 | 10.8 | 2.6 | 1,046 | 188 |
| 110,961 | 4,568 | 222 | 59.6 | 2.3 | 5,425 | 215 |

**B (server-side Cov2Tree)**

| # nodes | Naive algorithm compute time (ms) | Treenome algorithm compute time (ms) | Naive algorithm time to render (s) | Treenome algorithm time to render (s) | Naive algorithm web browser memory usage (MB) | Treenome algorithm web browser memory usage (MB) |
|---|---|---|---|---|---|---|
| 19,926 | 322 | 36 | 3.3 | 2.8 | 340 | 219 |
| 56,794 | 663 | 93 | 4.2 | 3.4 | 521 | 268 |
| 109,862 | 1,520 | 237 | 8.0 | 2.7 | 750 | 405 |

We quantified the performance of our phylogenetically-informed algorithm for drawing mutations by comparing it to a naive approach that does not fully utilize the phylogenetic relationship between samples. The naive algorithm enumerates the full set of mutations in each sample genome and displays each mutation as a distinct shape. We implemented both algorithms in **(A)** a version of Taxonium that performs all operations locally to the client, using the 9/21/22 global SARS-CoV-2 tree with 6.2M samples (public-2022-09-21.all.masked.jsonl in Table S2). We also implemented both algorithms in **(B)** a server-side version mirroring the behavior of cov2tree.org, using the Cov2Tree tree with 5.9M samples on 9/23/2022 (see Sanderson 2022 for a detailed description of the client-side and server-side modes). Each row contains results for a sparsified tree displayed at a different zoom level (see Implementation), with the final row of each table containing the tree displayed when fully zoomed out.

Compute time is the time taken to compute the data representing shapes to be drawn, measured with *performance.now()* in JavaScript. Time to render is the time elapsed between clicking “Enable Treenome Browser” and the mutations becoming visible, measured with a stopwatch. We computed memory usage with the *about:memory* tab in Firefox by taking measurements of Taxonium’s memory usage before and after Treenome Browser was enabled and subtracting the difference. We performed all tests on a MacBook Air with 16 GB of RAM.
